# Age-dependent Pavlovian biases influence motor decision-making

**DOI:** 10.1101/199331

**Authors:** Xiuli Chen, Robb B. Rutledge, Harriet R. Brown, Raymond J. Dolan, Sven Bestmann, Joseph M. Galea

**Affiliations:** School of Psychology, University of Birmingham, B15 2TT, UK; Max Planck University College London Centre for Computational Psychiatry and Ageing Research, London, WC1B 5EH, UK; Wellcome Centre for Human Neuroimaging, University College London, London, WC1N 3BG, UK; Sobell Department of Motor Neuroscience and Movement Disorders, UCL Institute of Neurology, University College London, London, WC1N 3BG, UK

**Author notes:** Corresponding authors; Xiuli Chen School of Psychology University of Birmingham B15 2TT, UK, Joseph Galea School of Psychology University of Birmingham B15 2TT, UK. Authors contributed equally (Sven Bestmann, Joseph M. Galea).

## Abstract

Healthy ageing is associated with decreased risk taking in motor^1^ and economic^2-4^ decision-making. However, it is unknown whether a single underlying mechanism explains these changes. Age-related changes in economic risk taking are explained by reduced Pavlovian biases that promote action toward reward^2, 5, 6^. Although Pavlovian biases also promote inaction in the face of punishment, the role such Pavlovian biases play in motor decision-making, which additionally depends on estimating the probability of successfully executing an action^7-10^, is unknown. To address this, we developed a novel app-based motor decision-making task to measure sensitivity to reward and punishment when subjects (n=26,532) made a ‘go/no-go’ motor gamble based on the perceived ability to execute a complex action. Using a newly established approach-avoidance computational model^2, 6^, we show motor decision-making is also subject to Pavlovian influences, and that healthy ageing is mainly associated with a reduction in Pavlovian bias toward reward. In a subset of participants playing an independent economic decision-making task (n=17,220), we demonstrate similar decision-making tendencies across motor and economic domains. Computational models that incorporate Pavlovian biases thus provide unifying accounts for motor and economic decision-making.

Optimal decision-making requires choices that maximise reward and minimise punishment^11^. To achieve this, humans rely on two key mechanisms; a flexible, instrumental, value-dependent process, and a hard-wired, Pavlovian, value-independent process^11-13^. Economic decision-making is often described using parametric decision models based on prospect theory that operationalise instrumental (value-dependent) concepts such as risk and loss aversion^14-17^. However, it has recently been shown that Pavlovian biases, which promote action towards reward and inaction in the face of punishment irrespective of option value^5, 11,18^, help to explain aberrant choice behaviour. For instance, the best explanation for the diminished economic risk-taking observed in older adults is a reduction in dopamine-dependent Pavlovian attraction to potential reward^2, 5^, suggesting that Pavlovian processes play a key role in explaining age-related changes in economic decision-making.

In contrast to economic decision-making, motor decision-making requires weighting potential rewards and punishments against the probability of successfully executing an action^7, 19-21^. Motor decision-making has primarily been explained in the context of instrumental-based processes^1, 7-10, 22^. Within this framework, older adults display reduced risk-seeking behaviour^1^. However, given recent findings in economic decision-making^2^, we asked whether Pavlovian biases might provide a more parsimonious explanation of age-related changes in motor decision-making. Although there is strong evidence that Pavlovian biases shape motor performance^23-26^, and that healthy ageing leads to a reduction in Pavlovian biases on motor performance^27, 28^, it is currently unknown whether Pavlovian biases influence motor decision-making. Sampling a large population through an app-based motor-decision game, we provide a novel demonstration that Pavlovian biases have a substantial impact on motor decisions, and are able to explain age-related changes in risk taking during motor decision-making.

We developed a novel app-based motor decision-making task that examined participant sensitivity to reward (gaining points) and punishment (losing points) when making a ‘go/no-go’ decision based on their perceived ability to successfully execute a motor action (Figure 1a, b). Using an app-based platform (‘How do you deal with pressure?’ The Great Brain Experiment: www.thegreatbrainexperiment.com)^18^, ^29, 30^, we obtained data from a large cohort (n=26,532; 15,911 males) in which six age groups were considered: 18-24yrs: n=5889; 25-29yrs: n=4705; 30-39yrs: n=7333; 40-49yrs: n=4834; 50-59yrs: n=2452; and 60+yrs: n=1319 (Figure 1c; see Supplementary Methods/ Figure S1)^18, 29, 30^.

**Figure 1:**
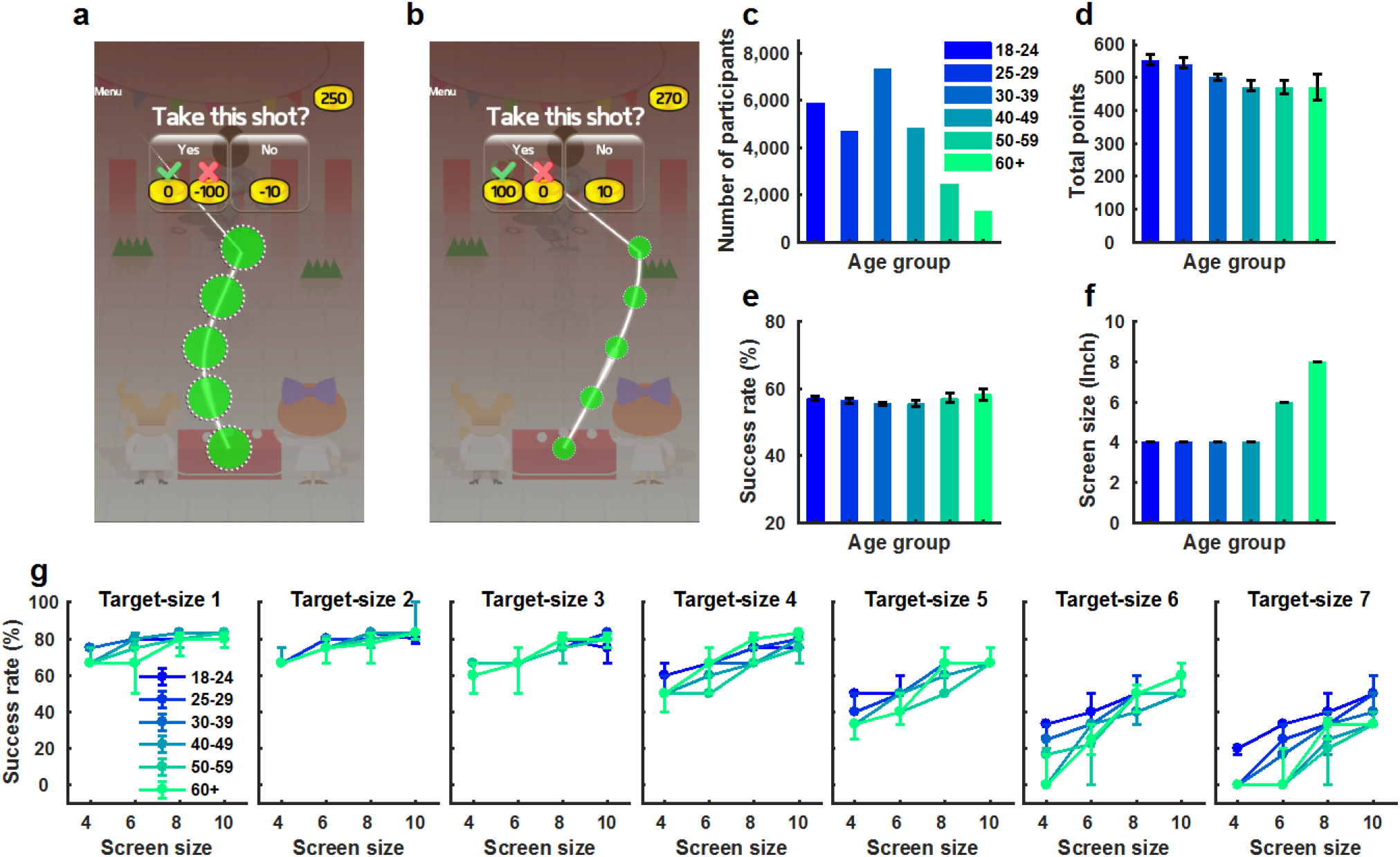
Motor gamble task and overall performance. (**a**) Game interface: an example of a punishment trial for target-size 1 (1: largest target size; 7: smallest target size); Participants decided whether to skip the tapping task and stick with a small punishment (-10 points) or gamble on successfully executing the action. If successful then they avoid the punishment (lose 0 points); otherwise, they received a greater punishment (-100 points); (**b**) A reward trial for target-size 7; (**c**) The number of participants in each age group; (**d**) Final points achieved across age groups; (**e**) The overall success rate (%) for executing the tapping action across age groups; (**f**) The screen size (inches) of the devices used across age groups; (**g**) Success rate (%) for executing the tapping action given the age, the screen size, and target-size (1: largest target size; 7: smallest target size). Bars/Dots and error bars represent medians and bootstrapped 95%CIs.

The game required participants to sequentially tap 5 targets distributed along a pre-defined path that could vary in both curvature and direction (Figure 1a, b; see Methods). If a participant accurately tapped all 5 targets successfully within 1.2 seconds, then the action was considered a success. There were 7 different target sizes, with the task becoming progressively more difficult as target size decreased (Figure 1a, b; see Methods). At the beginning of each trial, participants saw the required action and were asked whether they wanted to take the motor gamble. There were two types of trials: reward and punishment. For reward trials, participants had to decide whether to skip the trial and stick with a small reward (10 points) or gamble on successfully executing the tapping action (Figure 1b). If successful they received a greater reward (20, 60 or 100 points) or 0 points if they failed. For punishment trials, participants had to decide whether to skip the trial and stick with a small punishment (-10 points), or gamble on successfully executing the tapping action (Figure 1a). If successful, they lost nothing (lose 0 points) but failure resulted in a greater punishment (−20, −60 or −100 points). Participants began with 250 points and the overall goal was to accumulate as many points as possible. All trial-by-trial data (including tasks parameters, behavioural results, modelling results and accompanying code) are available on our open-access data depository (https://osf.io/fu9be/).

We found that older adults won fewer total points than younger adults (Figure 1d; r=-0.047, p<0.001; all *r* values represent a partial correlation between the measurement of interest and age, whilst controlling for the effects of gender and education; *p* values were computed by permutation test; see Methods). The final points accumulated during this task were dependent on (1) the decisions made (to gamble or not) and (2) the motor performance (success rate of executing the tapping action). Therefore, prior to examining participant choice behaviour it was crucial to determine whether motor performance differed across age groups.

Although success on the motor task was similar across age groups (Figure 1e, r=0.006, p=0.329), older adults used devices with larger screen sizes than younger age groups (Figure 1f, r=0.279, p<0.001). As target size was scaled to device screen size (see Methods), we assessed how the relationship between age, target size and screen size affected motor performance. We found that decreased success rate was linked to a combination of smaller target sizes, smaller screen sizes and older age (Figure 1g, stepwise regression winning model: success rate =1 - 0.003*age*target size + 0.002*age*screen size + 0.005*target size*screen size; all p<0.001; Adjusted R^2^=0.213). Therefore, we next assessed choice behaviour in the context of how these factors influenced motor performance on a trial-by-trial basis.

Participants were asked to make decisions between a gamble option and a certain option. Each option can be characterised by its potential outcomes, weighted by the probability of each outcome (i.e. Expected Value^31^). For the gamble option, the expected value is given by: EV_gamble_=P_success_V_success_+(1-P_success_)V_failed_, where P_success_ is the probability of successfully executing the tapping action; V_success_ is the points received if successful; V_failed_ is the points received on failure. The expected value of the certain option (EV_certain_) is V_certain_ and the probability of receiving this value is 1. We calculated P_success_ by estimating the probability of motor success based on a participant’s age, screen size of the device used and target-size level (Figure 1g; see Methods). By comparing choice behaviour given the difference between these two options (EV_gamble_-EV_certain_), we were then able to examine the influence of ageing on motor decisions while controlling for differences in motor performance due to age, screen size and target size. However, this formulation relied on an assumption that participants had a good estimate of their probability of success. To test whether this was true, we recruited an additional 60 participants (10 in each age group) who were asked to estimate their probability of success (from 0% to 100% in steps of 10%; see Methods) after being shown the target size and trajectory. After this estimate, they were then asked to perform the tapping action (whilst ignoring the decision-making part of the game). Similar to previous work^1, 7, 20^, we found participants were able to reliably estimate the probability of motor success (Figure 2a), and this estimate did not differ across age groups (Figure 2b; one-way ANOVA: F_(5,54)_=0.859, p=0.515).

**Figure 2:**
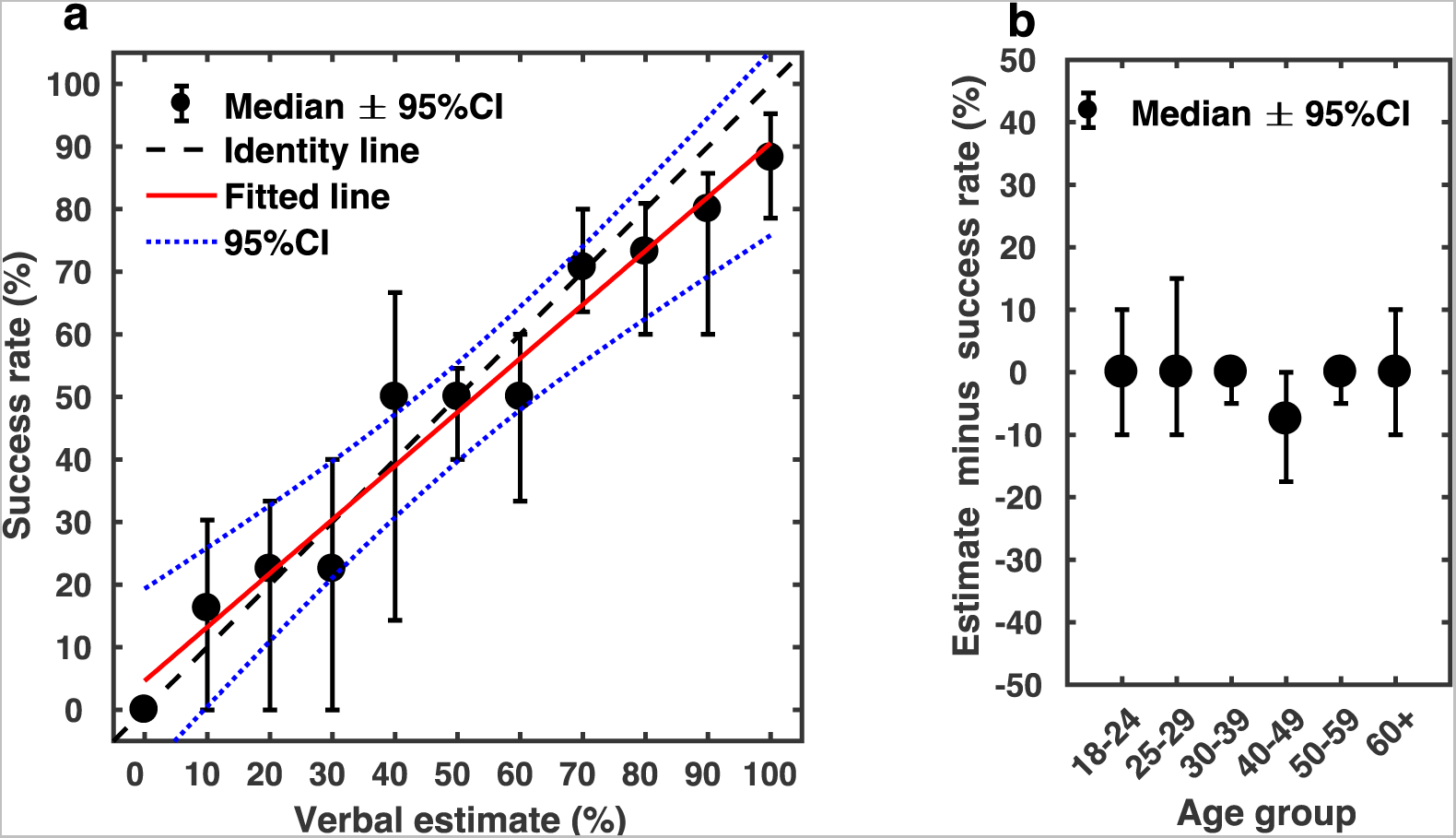
Participant ability to estimate motor performance success. (**a**) For each participant (n=60; 10 in each group), we calculated an average success rate for each available verbal estimate value (0% to 100% with 10% increment). Each black dot represents the median success rate (y-axis) across participants who gave that certain verbal estimate value (x-axis), and error bars represent bootstrapped 95% CI across participants; (**b**) The estimation error for each age group. For each participant, estimation error was calculated as the median error (on each trial: estimate % - 100% if successful, 0% if failed) across all trials. Black dots and error bars represent the medians and bootstrapped 95%CIs.

We found a significant decrease in the proportion of trials in which participants chose to gamble across the lifespan in reward trials (Figure 3a; r=-0.190; p<0.001), and to a lesser extent in punishment trials (Figure 3b; r=-0.052; p<0.001). To understand these results, age-related changes in choice behaviour had to be examined given the difference between these two value options (EV_gamble_-EV_certain_). Interestingly, in reward trials, there was a gradual and monotonic decrease in gamble rate across the lifespan which appeared independent of the EV_gamble_-EV_certain_ value (right side of Figure 3c). In contrast, for punishment trials, older adults displayed a higher gamble rate during high risk gambles (e.g., EV_gamble_-EV_certain_=-90), but conversely a reduced gamble rate during low risk gambles (e.g., Figure 3c; EV_gamble_-EV_certain_=0).

**Figure 3:**
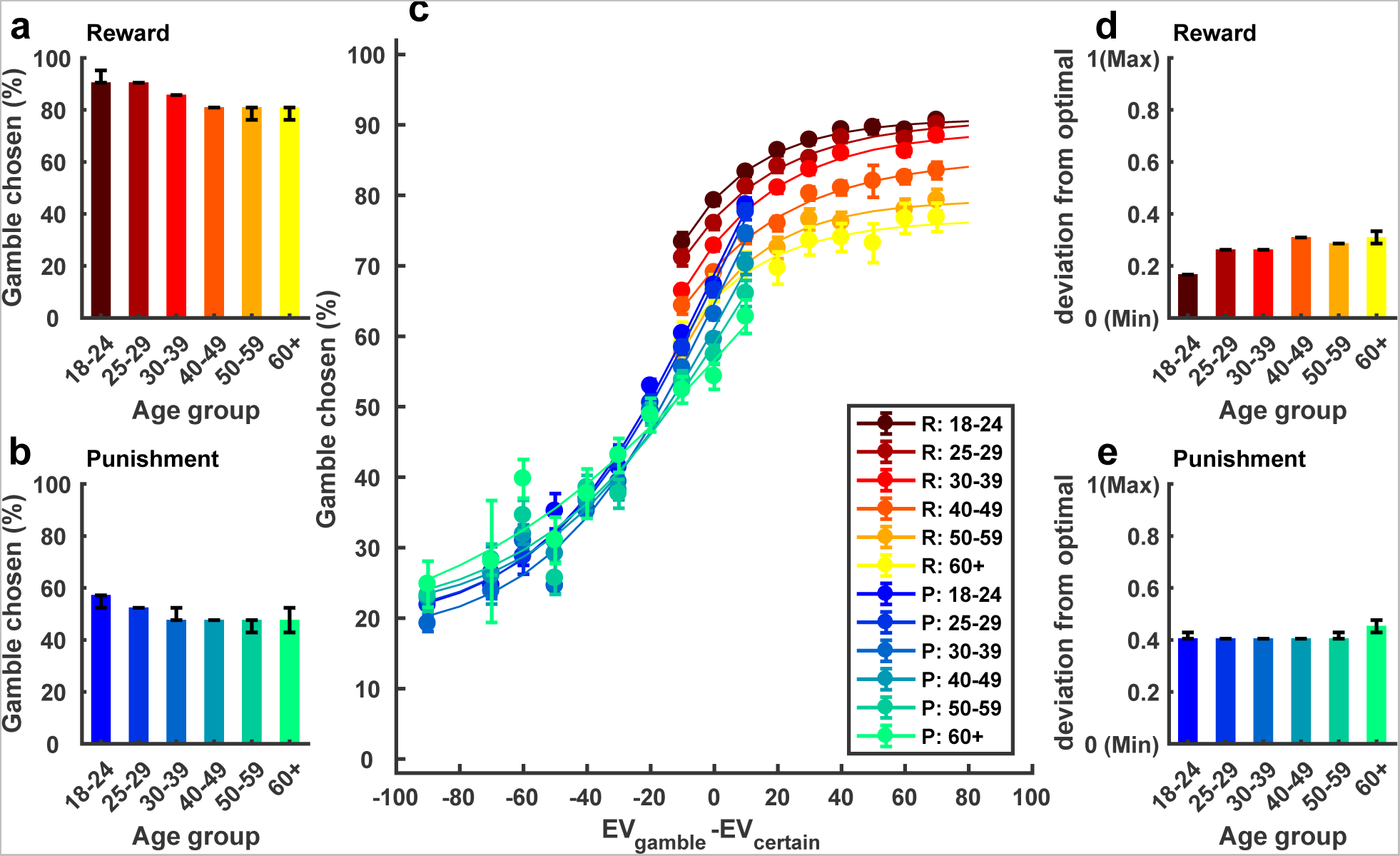
The proportion (%) of trials in which participants chose to gamble. (**a**) Gamble rate in the reward and **(b)** punishment domain; (**c**) Propensity to choose the gamble option as a function of EV_gamble_ – EV_certain_ (data was grouped into bin sizes of 10). As indicated in the legend, each of the warm colours represents one age group in the reward (R) condition, and each of the cool colours represents one age group in the punishment (P) condition. The lines are fitted lines to y=a*exp(-b*x)+c; R^2^=0.979 **±** 0.022; **(d)** Discrepancy between choice behaviour and optimal decisions in the reward domain. Specifically, using EV_gamble_-EV_certain_ we calculated whether the optimal decision on each trial was to gamble (1) or skip (0). We then subtracted this value from the observed behaviour of the participant (gamble=1, skip =0). If the average absolute difference between these values across trials was 0, then a participant was deemed as an optimal decision-maker; **(e)** Discrepancy between choice behaviour and optimal decisions in the punishment domain. Bars and error bars represent medians across the participants and bootstrap 95% CIs.

Given these results, do older adults make less optimal motor decisions? An ideal (optimal) decision-maker chooses the option that has the higher expected value, and we therefore compared participant’s choice behaviour with the optimal behaviour. Specifically, using EV_gamble_-EV_certain_ we calculated whether the optimal decision on each trial was to gamble or decline (coded 1 and 0 respectively). We then subtracted this value from the observed behaviour of the participant (also coded gamble = 1, decline = 0). If the average absolute difference between these values across trials was 0, then a participant was deemed an optimal decision-maker. In reward trials, there was progressive deviation from optimality across the lifespan (Figure 3d; r=0.232; p<0.001). In contrast, for punishment trials, all age groups showed a similar level of sub-optimality (Figure 3e; r=0; p=0.999). Therefore, the most pronounced effect of ageing on motor decision-making was a value-independent decrease in gamble rate during reward trials which led to a significant decrease in optimality.

While these data portray many similarities with decision-making under risk^14, 15^, there are also clear differences. For example, decision-making models based on prospect theory are not able to explain the gradual, monotonic and value independent decrease in gamble rate across the life span observed during the reward trials^5^ (Figure S1). We predicted that such dichotomies represented the contribution of value-independent Pavlovian approach-avoidance biases to motor decision-making behaviour. To test this prediction, we modelled the choice behaviour using an established decision-making model based on prospect theory, and a newly introduced model which included Pavlovian approach-avoidance parameters^2, 5^ (see Methods). The prospect theory model included three components: (1) loss aversion parameter (λ) (2) risk preference parameter (*α*) and (3) stochasticity of decision-making captured by an inverse temperature parameter (*μ*). The loss aversion coefficient (*λ*) represents the relative (multiplicative) weighting of losses relative to gains, which was set to 1 as there were no gambles with both positive and negative outcomes in our task. The risk preference parameter (*α*) represents the diminishing sensitivity to change in value with an increase in absolute value (value-dependent). The logit parameter *μ*.is the sensitivity of the choice probability to an option value difference. In addition to these parameters, the Pavlovian approach-avoidance model included value-independent parameters exclusively for reward (*δ*^*+.*^).and punishment (*δ*^*-.*^) trials. Positive or negative values of these parameters correspond, respectively, to an increased or decreased probability of gambling without regard to the value of gamble (see Methods; Eq 3). We found an approach-avoidance decision model with 4 parameters (a single risk preference parameter: *α*, the inverse temperature parameter: *μ*, value-independent parameters exclusively for reward *δ*^*+.*^and punishment *δ*^*-.*^trials) fitted the motor gamble (choice) data better than any decision model based on prospect theory (Table S1, Figure S2 & S3; see Methods for model comparison).

Using this preferred model, we observed age-related changes across the reward and punishment domains for both value-dependent and independent parameters. However, the most striking effect was a large decrease in Pavlovian attraction which facilitates action in pursuit of reward. Specifically, we found that healthy ageing did not affect the stochasticity parameter, *μ* (r=-0.001, p=0.871), but was associated with a decrease in the risk preference parameter, *α* (Figure 4a,d; r=-0.105, p<0.001). The winning model included a single α parameter, which represented different value-dependent biases in reward and punishment (*α*<1 indicated risk aversion in reward domain and *α*<1 represented risk-seeking in punishment domain; *α*=1 represented risk-neutral; see Supplementary Methods). Therefore, older adults displayed a similar increase in value-dependent biases across the reward and punishment domain. The greater risk-seeking effect in the punishment domain was offset by the fact that ageing was also linked with greater Pavlovian avoidance (Figure 4b, d; *δ*^*-.*^; r=-0.076, p<0.001), an effect not previously observed in economic decision-making^2^. Such opponent effects between value-dependent and value-independent parameters help to explain the complex changes observed with ageing during punishment trials (Figure 3c). Nevertheless, the largest effect of ageing was a substantial decrease in Pavlovian attraction (Figure 4b, d; *δ*^*+*^; r=-0.146, p<0.001). We found a similar decline for both sexes (Figure S4; male: n=15911, r=-0.140, p<0.001; female: n=10621, r=-0.143, p<0.001), and across all education levels (Figure S5; school: n=9171, r=-0.152, p<0.001; university: n=11281, r=- 0.142, p<0.001; advanced: n=6080, r=-0.136, p<0.001). Importantly, we did not observe this age-related effect for the temperature parameter (*μ*), indicating the changes observed in the risk and Pavlovian parameters were not simply a result of large participant numbers (Figure 4d).

**Figure 4:**
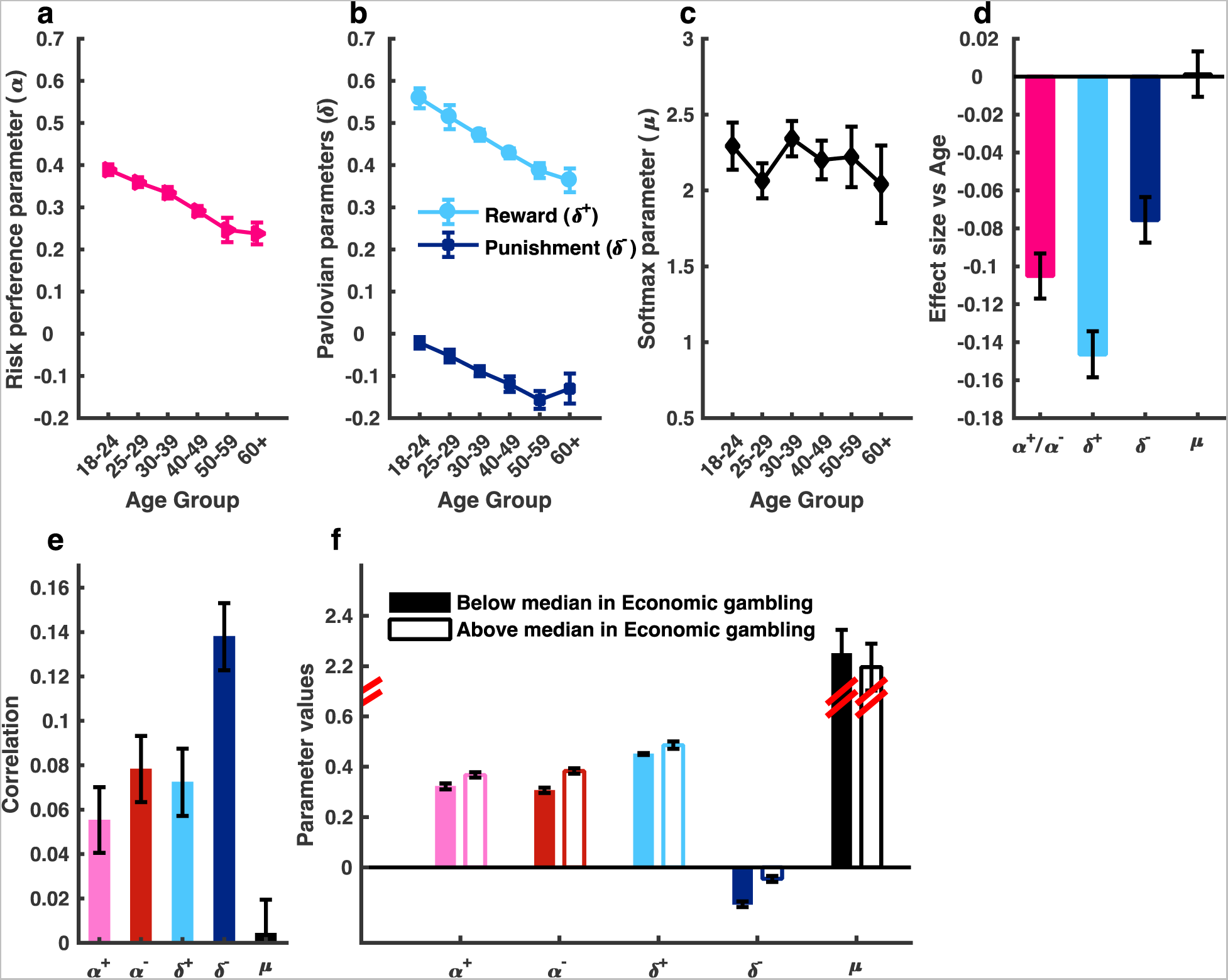
The change in approach-avoidance model parameters across the life span, and the relationship between approach-avoidance model parameters across a motor and economic gambling task. (**a**) α across age groups; (**b**) δ^*-.*^and δ^*+.*^across age groups; (**c**) μ across age groups; (**d**) Age-related decline across the reward and punishment domain. The largest effect size was observed for the Pavlovian approach parameter (δ^*+.*^). This age-related effect was not observed for the temperature parameter, *μ*; (**e**) Positive correlation across independent motor and economic decision tasks for the main approach-avoidance model parameters. Note, the single α parameter of the motor decision-making model was correlated with both the α^-^ and α^+^ parameters of the decision-making model. This positive correlation was not observed for the temperature parameter, *μ*; (**f**) Motor decision-making approach-avoidance parameter values median split by economic parameter values. Filled bars denote participants with below-median values in the economic gambling task; Hollow bars for above-median. The participants with above-median risk parameters and Pavlovian parameters in the economic decision task had higher risk parameters and Pavlovian parameters in the motor gambling task. This median split effect was not observed for the temperature parameter, *μ*; Bars/error bars reflect medians/bootstrapped 95%CIs.

Finally, through this app-based platform a subset of participants (n=17,220) also performed an economic decision-making gambling task in which a similar approach-avoidance model was used to explain choice behaviour^2^. Through correlation and median-split analysis we found a significant positive relationship for all main model parameters between the tasks (Figure 4e, f), consistent with participants manifesting similar decision-making tendencies across motor and economic domains. This relationship was relatively consistent across the lifespan whereby a positive correlation existed between these parameters within each age group (Figure S6, S7). Once again, we did not observe this correlation for the temperature parameter (*μ*).

Making decisions under uncertainty is crucial in everyday life, whether it is managing retirement funds, choosing a career, or deciding between pulling out or not on to a busy road whilst driving. The latter example describes motor decision-making, a unique kind of decision which requires weighting potential rewards and punishments against the probability of successfully executing an action^7, 19-21^, and often with immediate outcomes. Although healthy ageing has been associated with decreased risk taking across both motor^1^ and economic^2-4^ decision-making, it is heretofore unknown whether a single underlying mechanism might explain these changes. We addressed this question using a novel motor gambling task that exploited an app-based platform which enabled us to collect a large cohort of data. Unlike previous work on motor decision-making^7, 19-21^, we considered choice behaviour in relation to both value-dependent instrumental and value-independent Pavlovian processes^5, 11, 18^. We found age-related changes across the punishment and reward domain for both value-dependent and independent parameters. However, the most striking effect of ageing was a decrease in Pavlovian attraction which facilitates action in pursuit of reward. Through this app-based platform, we compared a subset of participant’s choice behaviour during motor and economic decision-making^2^ and found similar decision-making tendencies across motor and economic domains.

Our large cohort and use of a newly established approach-avoidance computational model^2, 6^ enabled us to detect subtle age-related changes in choice behaviour and surprising interactions between value-independent and value-dependent processes. For instance, the risk aversion parameter (*α*: instrumental value-dependent process) was on average less than 1 across all age groups, indicating risk aversion in reward, and risk seeking in punishment. This choice behaviour was best explained using a single parameter, signifying a similar degree of risk aversion in reward and risk-seeking in punishment. Importantly, this value progressively decreased with age suggesting that older adults showed similar increased risk-aversion for reward and risk-seeking for punishment. This is in line with previous economic decision-making work which revealed older adults weigh certainty (achieving the small reward or avoiding the small punishment) more heavily than younger adults^32^. Interestingly, the greater risk-seeking in the punishment domain was offset by the fact that ageing also led to greater Pavlovian avoidance, an effect not observed in economic decision-making^2^. It is the interaction between value-dependent and independent parameters that help explain not only the complex changes observed with ageing during punishment trials but also the lack of difference across age groups for punishment-based optimality. Crucially, previous work in motor decision-making using only instrumental-based processes would not have detected such complex behavioural interactions^1, 7, 21^. The underlying mechanism behind age-dependent increases in Pavlovian avoidance is unknown. It has been suggested that the neurobiology behind Pavlovian avoidance may involve opponency between the dopaminergic and serotonergic systems^33^. Despite there being evidence of age-related decline in serotonin receptor availability^34^, it remains an open question as to the link between serotonin and Pavlovian avoidance during either motor or economic decision-making.

The strongest effect of ageing was a decrease in Pavlovian attraction to reward. As all age groups displayed risk aversion during reward trials, this decrease in Pavlovian attraction led to greater sub-optimality in older adults. These results are strikingly similar to the ones observed in economic decision-making^2^, suggesting Pavlovian attraction plays a pivotal role in explaining age-related changes to reward across both motor and economic decision-making. During economic decision-making, it has recently been shown that boosting dopamine with L-DOPA increases the influence of Pavlovian attraction on choice behaviour^5^. In addition, healthy ageing is associated with a gradual decline in dopamine availability^35, 36^ and neural responses to reward^37^. Therefore, it is possible that the decrease in Pavlovian attraction during motor decision-making in older adults is a result of an age-dependent decrease in dopamine availability.

More broadly, the current work shows the importance of both instrumental value-dependent and Pavlovian value-independent processes on motor decision-making. However, task design may play an important role in determining the size of Pavlovian influences. Here we used a ‘go/no-go’ decision-making task as previous literature has shown the ‘go/no-go’ component induces strong Pavlovian biases^11, 25, 38^. It remains to be seen whether computational models including Pavlovian biases provide a better description of choice behaviour during other motor decision-making tasks which do not involve a ‘go/no-go’ component^1, 7-10, 22^.

Finally, participants showed similar decision-making tendencies for both instrumental (value-dependent) and Pavlovian (value-independent) parameters across motor and economic domains. This extends previous work that revealed a similar relationship with parameters derived from parametric decision models based on prospect theory^7, 10^, and reinforces the view that the mechanisms which control cognitive (economic) and motor decision-making are integrated^39^. However, the correlation between the tasks was small, around r=0.1, suggesting that while participants showed similar behavioural trends across the two tasks, their performance in motor and economic domains was also distinct. Interestingly, the approach-avoidance model not only fitted choice data substantially better for the motor decision-making task, relative to the economic task, but the effect size relating to age was also nearly double in size for all parameters^2^. This indicates that while there are clear similarities between cognitive and motor decision-making, computational models including Pavlovian biases appear to be particularly important for explaining motor decision-making.

In conclusion, Pavlovian biases play an important role in not only explaining motor decision-making behaviour but also the changes which occur through normal ageing. This provides a greater understanding of the processes which shape motor decision-making across the lifespan, and may afford essential information for developing population wide translational interventions such as promoting activity in older adults.

## Methods

### Participants

We tested 26,532 participants (15,911 males, aged 18-70+) who completed the task between November 20, 2013 and August 15, 2015. Data were only included if users fully completed the game and it was their first attempt. We also recruited an additional 60 participants (29 males, aged 18-70+) who were asked to estimate their success rate (motor performance). All participants gave informed consent and the Research Ethics Committee of University College London approved the study.

### Materials and apparatus

Using an app-based platform (The Great Brain Experiment: www.thegreatbrainexperiment.com) we developed a motor decision-making task (‘How do I deal with pressure?’) which is freely available for Apple iOS and Google Android systems. The game runs in a 640x960 (3:4 ratio) pixel area, which is then scaled to fit the screen whist maintaining this ratio. The game required participants to ‘throw’ a ball at a coconut in an attempt to knock it off its perch. This was achieved by tapping 5 sequential targets along a pre-defined path. The path was characterised by an angle parameter that represented a section of a sine curve, in degrees. The curves were drawn from the bottom (the starting point) to top of the game window (Figure 1a, b). For example, if the angle parameter was 360, then one complete cycle of the sine curve was used to draw the curve. During the task, the angle was randomly chosen between 0 and 360. The 5 targets were evenly spaced along the curves. If the participant tapped all 5 targets sequentially (from bottom to top) within 1.2 seconds, then the action was considered a success and the coconut was hit. If the participant failed to tap all 5 targets accurately or within the allotted time then the action was considered a failure and the ball sailed past the coconut. In addition, the action was deemed a failure if participants did not start the tapping action within 7 seconds after they chosen to do the tapping. There were 7 different target sizes across trials with the tapping action becoming more difficult as the target size was reduced. However, as mentioned above, the game interface was scaled to screen size. Therefore, motor performance (success rate) was examined relative to the interaction between target size and screen size (Figure 1g). All trial-by-trial data (including tasks parameters, behavioural results, modelling results and accompanying code) are available on our open-access data depository (https://osf.io/fu9be/).

### Motor gambling task

At the beginning of each trial, participants were shown the action required (i.e. the position and size of the 5 targets) and were asked to make a motor gamble. There were two types of trials: reward trials and punishment trials (Figure 1a, b). For reward trials, participants had to decide whether to skip the trial and stick with a small reward (10 points) or gamble on successfully executing the ‘throw’. If successful they received a greater reward (20, 60 or 100 points) but 0 points if they failed. For punishment trials, participants had to decide whether to skip the trial and stick with a small punishment (-10 points) or gamble on successfully executing the ‘throw’. If successful they lost nothing (0 points) but failure resulted in a greater punishment (−20, −60 or −100 points). Hence, there were 6 value combinations. Each combination was repeated for each of the 7 different target sizes (6 values x 7 target sizes = 42 trials). Although there were 7 blocks of the game this did not directly relate to the 7 target sizes. In order to maintain a level of unpredictability, the first 3 blocks included random presentation of the 3 largest (easiest) target sizes, the next 3 blocks included target sizes 4-6 and the final block included the smallest (most difficult) target size. Participants began with 250 points and the overall goal was to accumulate as many points as possible.

For the control study (Figure 2) which examined participant’s ability to estimate their probability of success, individuals were asked to estimate their probability of motor success (0% to 100% in steps of 10%) after being shown the target size and trajectory. After this estimate, they were then asked to perform the tapping action, whilst ignoring the decision-making part of the game.

### Data analysis

Matlab (Mathworks, USA) was used for all data analysis. We reported partial correlation coefficients (r) for the relationships between task measures and age, whilst controlling for the effects of gender and education. All p values were computed based on permutation tests using 100,000 random shuffles of age labels to determine null distributions^2^. Bootstrapped 95% confidence intervals were computed based on 100,000 resamples with replacement in each age group^2^.

### Parametric models

On each trial participants faced a gamble that contained a certain option (CO) involving a payoff of certain points (+10 in reward trials and −10 in punishment trials), and a gambling option (GO) in which the outcome depended on a probability of successfully executing the tapping action. The probability was estimated given a participant’s age, screen size of the device used and target-size level (Figure 1g). Specifically, the probability of success for a participant within a certain age group, using a certain screen size and facing a certain target size on each trial was estimated using the average success rate across all the participants with the same age, same screen size, and facing the same target size. Given the small amount of trials we had for each participant at each target size to estimate their probability of success, we believed this group average approach was the most valid estimate of success probability. However, we also conducted the analysis when success probability was estimated based on each individual’s own data (i.e. the probability of success for a participant facing a certain target size was estimated using their own success rate over the same target size). Importantly, our findings still hold (Figure S8). We modelled participant motor gamble choices using an established decision-making model based on prospect theory^15^ and a newly introduced model which included an extra Pavlovian approach-avoidance^2, 5, 6^ component. In the following, we first describe the prospect theory models, followed by the approach-avoidance models.

*Parametric decision-making model based on prospect theory:* There are three key components in prospect theory models. The first component is the value function. According to prospect theory, the subjective desirability of outcomes is modelled as transformations of objective task quantities. The subjective desirability of the outcomes, *O* (the points in this case) was modelled by a value function (2-part power function) of the form:

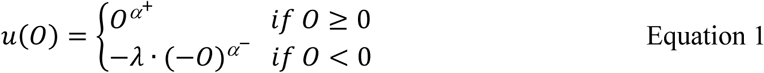

where, the risk preference parameter (*α*) represents the diminishing sensitivity to changes in values as the absolute value increases (if *α>1.*). The loss aversion coefficient (*λ*) represents the weighting of losses relative to gains, which was set to 1 as we did not have gambles with both positive and negative outcomes. The second component of a prospect theory model is the probability weighting function. Most prospect theory models assume that probabilities are weighted non-linearly. However, we found that the probability weighting parameter (*γ*) did not significantly improve the model fit (Table S1, we used a 1-parameter probability weighting function^40^: *w.*(*p*).*exp*(*−*(*−ln*(*p*)*γ*)).). Hence, probabilities and utilities were combined linearly in the form: *U.(p,0)=p*v(0)*. The third component of a prospect theory model is the choice function. The probability of choosing to gamble is given by the logit or soft-max function:

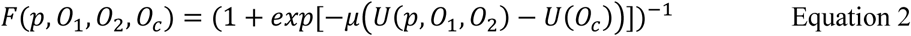

where *0*_*1*_.and *0*_*2*_.are the outcomes in the gamble option *[p.→ 0*_*1*_; *(1-p)→ 0*_*2*_], and *O*_*c*_ is the outcome of the certain option. The logit parameter *μ* is the sensitivity of the choice probability to the utility difference. In summary, our prospect theory models included the following free parameters: risk preference parameter (*α)* and stochasticity of decision-making according to the inverse temperature parameter (*μ*).

*Parametric approach-avoidance decision model:* Approach-avoidance models were based on the prospect theory models, but with an additional component that allows for value-independent influences to choose or not choose gambles. Specifically, Pavlovian parameter (*δ*).were added to the probability of choosing to gamble (Equation 2) as follows:

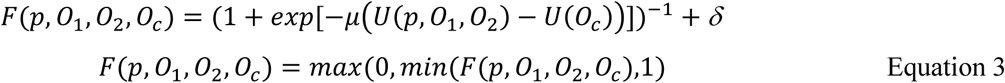

Positive or negative values of the parameter (*δ*).correspond respectively to an increased or decreased probability of gambling without regard to the value of gamble. Other parts of the models were identical to the prospect theory models. In summary, the approach-avoidance model included the following free parameters: risk preference parameter (*α*)., stochasticity of decision-making according to the inverse temperature parameter (*μ*) and Pavlovian parameter (*δ*).

### Parameter optimisation and model selection procedures

The models were fit to individual choice data. The method of maximum likelihood was used to estimate (fminsearch in Matlab) the parameter vector Θ given the participant choice (*y*) on each trial *(p, 0*_*1*_, *0*_*2*_, *0*_*c*_):

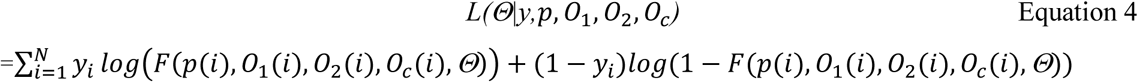

Where, *i* indexes the trial number; *N* is the number of trials; *y*_*i*_ indicates participant choice on trial *i*; *Θ* indicates the parameter vector to be estimated; (*p, 0*_*1*_, *0*_*2*_, *0*_*c*_).represent the gamble options on each trial. Parameters were constrained to the following ranges:*α:0→1; μ:0→10;δ:-1→1*The model was fit to each participant’s data, and the fitting was repeated at 200 random seed locations to avoid local minima.

For each key parameter of prospect theory and approach-avoidance models, we explored the possibility of using separate and single parameters for reward and punishment domains as well as a weighted or linear probability function. Therefore, we fitted each participant’s choice data with 24 models (Table S1). We used Akaike’s information criterion (AIC)^41^ and Bayesian information criterion (BIC)^42^ to compare model fits. Both of these represent a trade-off between the goodness of fit and complexity of the model and thus can guide optimal model selection.*pseudo r*^*2*^ was calculated with the null model in which *α μ* and *δ* were restricted to 0 (*pseudo* 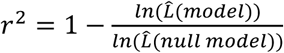, where 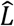 = Estimated likelihood).

## Competing interests

The authors declare no competing interests.

## Acknowledgements

We thank N. Millstone and Peter Zeidman for developing the app, R. Davis, C. Freemantle and R. Maddock for supporting data collection, D. Jackson, C. Brierley and C. Sheppard for supporting app development and R.C Miall for helpful comments on the manuscript. This work was supported by European Research Council Starter Grants MotMotLearn (637488) to JMG, ActSelectContext (260424) to SB. A Wellcome Trust Engaging Science: Brain Awareness Week Award (101252/Z/13/Z) supported app development. R.J.D is supported by the Wellcome Trust (078865/Z/05/Z). RR is supported by an MRC Career Development Award (MR/N02401X/1).

## Supplementary Methods

### Participants

**Figure S1:**
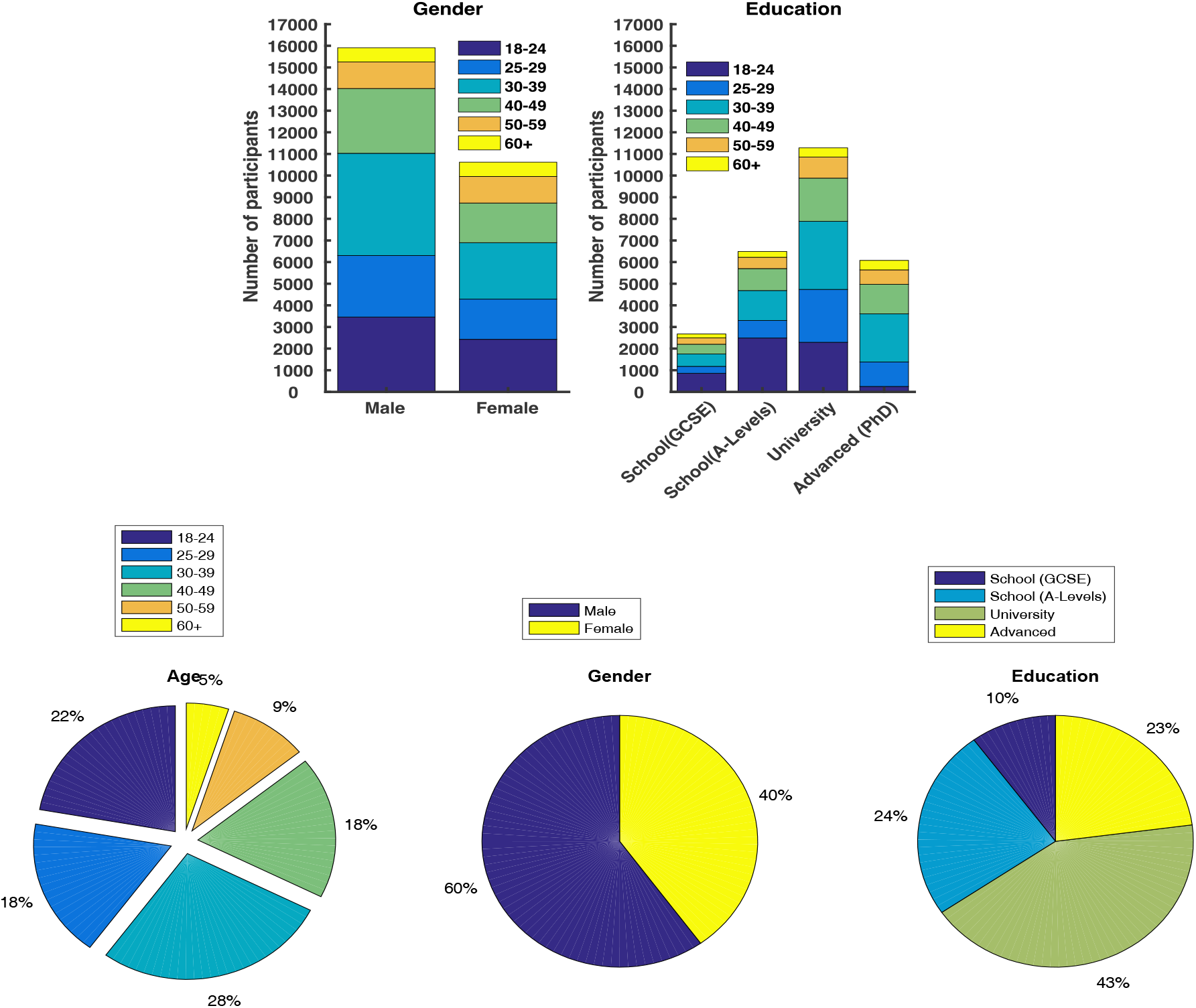
Additional participant demographics.

### Parametric decision-making model based on prospect theory

As mentioned in the Methods, the subjective desirability of outcomes was modelled as transformations of objective task quantities. The subjective desirability of the outcomes *O* (the points in this case) was modelled by a value function (2-part power function) of the form as in Equation 1. The risk preference parameter (*α*) represents the diminishing sensitivity to changes in values as the absolute value increases (if *α<1*). The risk preference parameter (*α<1*) is equivalent to risk aversion in the reward domain and risk seeking in the punishment domain, as demonstrated by the following examples. Imagine a gamble between a probabilistic reward: 50% of £20; 50% of £0 and a sure reward of £10. The objective expected value of the gamble is £10, similar to the certain option. Hence a risk neutral person would be indifferent between these two options. If *α=0.8*, the gamble would have a subjective value of 5.49, and the certain option would have a subjective value of 6.31, which results in participants being more likely to choose the certain option (i.e., risk aversion). Now imagine a gamble between a probabilistic punishment 50% of -£20; 50% of £0 and a sure punishment of -£10. The objective expected value of gamble is -£10, similar to the certain punishment option. If *α=0.8*, the gamble would have a subjective value of −5.49, and the certain option would have a subjective value of −6.31, which results in participants being more likely to choose the gamble option (i.e., risk seeking).

### Parametric approach-avoidance decision model

Approach-avoidance Models were based on the prospect theory models, but with an additional component that allows for value-independent influences to choose or not choose gambles i.e., Pavlovian parameters (*δ*). Positive or negative values of the parameter (*δ*).correspond respectively to an increased or decreased probability of gambling without regard to the value of gamble. Other parts of the models were identical to the prospect theory models.

## Supplementary Results

### Model parameter optimization and model selection

For each key parameter of prospect theory and approach-avoidance models, we explored the possibility of using separate and single parameters for gain and loss domains as well as a weighted or fixed probability function. Therefore, we fitted each participant’s choice data with 24 models (Table S1). We used Akaike’s information criterion (AIC) and Bayesian information criterion (BIC) to compare model fits (see main paper for relevant references). Both of these represent a trade-off between the goodness of fit and complexity of the model and thus can guide optimal model selection *pseudo r*^2^ was calculated with the null model in which *α,μ* and *δ* were restricted to 0 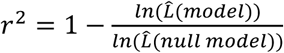, where 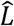 = Estimated likelihood). The preferred model’s behavioural predictions among both the prospect theory models ([*α*^*+*^,.*α*^*−*^, *μ*^*+*^, *μ*^*−*^.]; ID=4 Table S1) and the approach-avoidance models *([α, μ, δ*^*+*^, *δ*^*-.*^]; ID=10 Table S1) are plotted in Figure S2 and Figure S3, respectively.

**Table S1:**
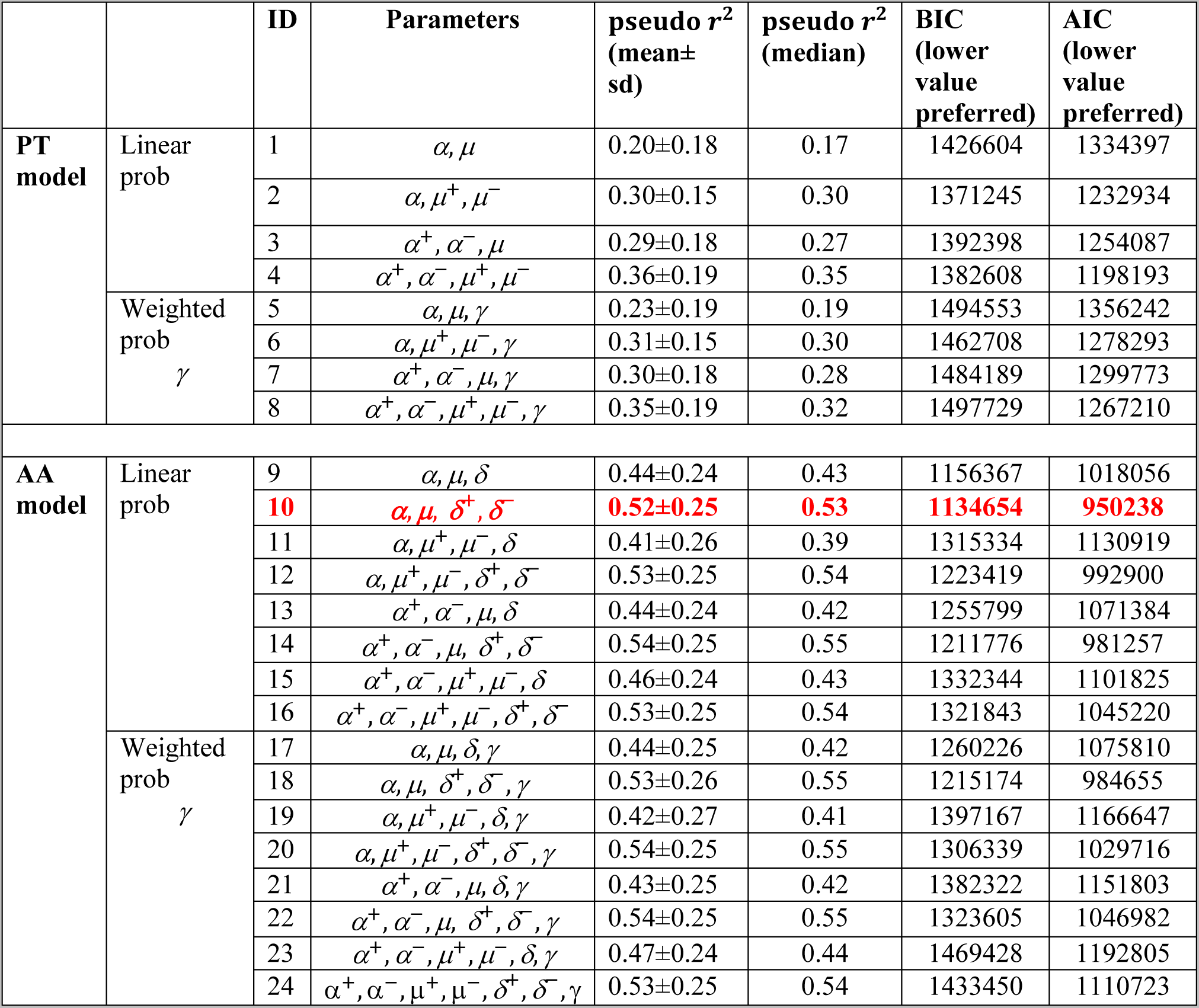
Comparison of decision-making models. The main parameters were (1) value function parameter (*α*); (2) the probability weighting parameter (*γ*); the Softmax temperature parameter (*μ*); the Pavlovian parameter (*δ*). For each key parameter of the prospect theory (PT) and approach-avoidance (AA) models, we explored the possibility of using separate and single parameters for reward and punishment domains as well as a weighted or fixed probability function (see Methods). According to AIC and BIC model comparison an approach-avoidance decision model (red; ID=10) fitted the choice (gamble) data better than established decision models based on prospect theory.

**Figure S2:**
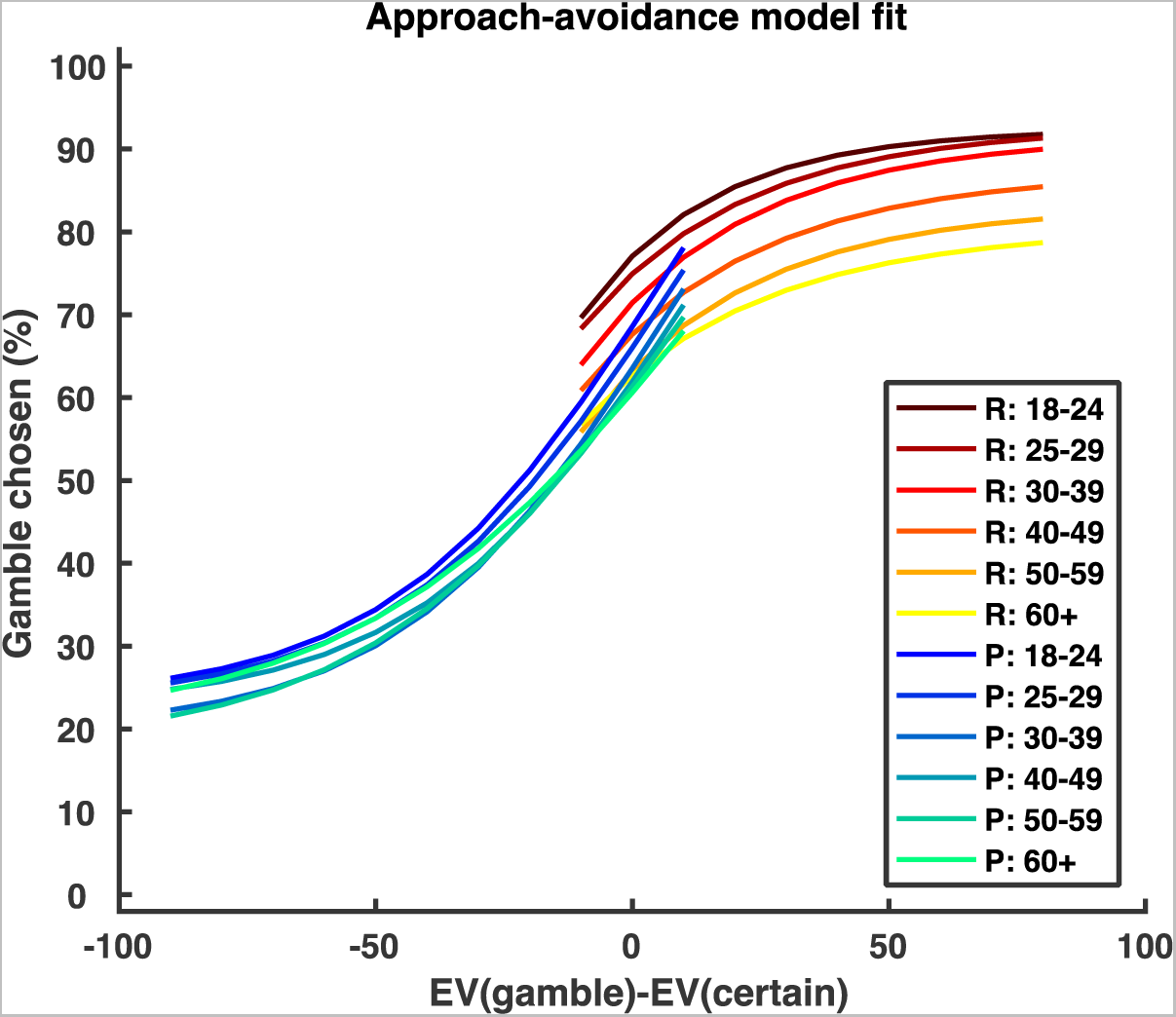
Average model fit across participants for the winning prospect theory model (ID=4 Table S1 [*α*^*+*^, *α*^*−*^, *μ*^*+*^, *μ*^*−*^]). The model cannot account for the observed differences in choice behaviour during the motor decision-making task across the lifespan, including (1) the value-independent differences across age groups in the reward domain (Figure 3); (2) the changes in gamble propensity observed in the punishment domain across age groups (the model fit shows almost no difference across the age groups in the punishment domain); (3) the higher gamble rate in the reward domain relative to the punishment domain when the difference between expected values [EV_gamble_-EV_certain_] is close to 0 (the model fit shows the opposite).

**Figure S3:**
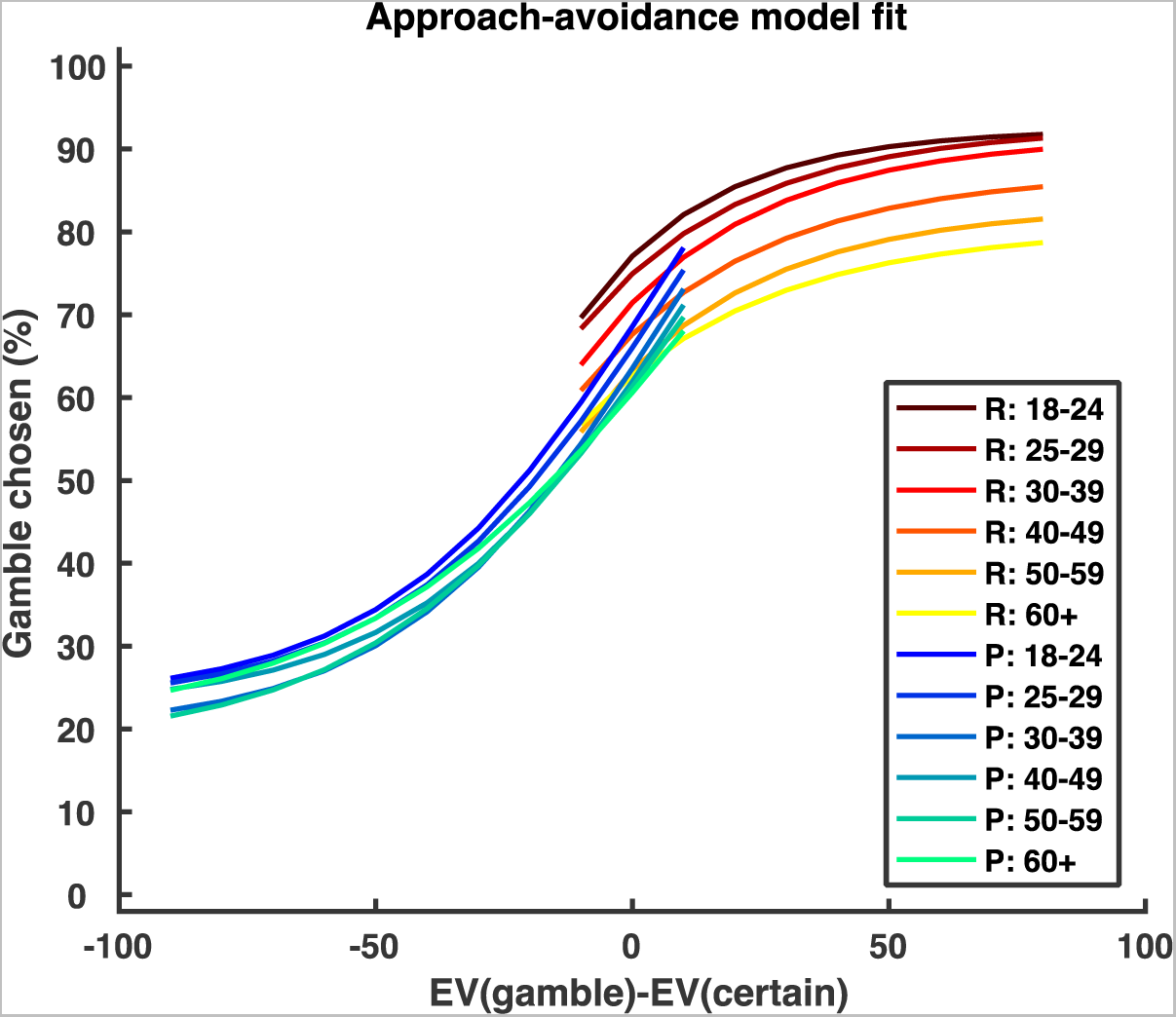
Average model fit across participants for the winning approach-avoidance model (ID=10 Table S1, *[α, μ, δ*^*+*^, *δ*^*−*^.]). The model does a far superior job of fitting choice behaviour during the motor decision-making task across the lifespan (Figure 2).s

### The aging effect for each gender and education level

We found similar aging effects in both males (n=15911, r=-0.140, p<0.001, Figure S) and females (n=10621, r=-0.143, p<0.001, Figure S), and across all education levels (school: n=9171, r=-0.152, p<0.001; university: n=11281, r=-0.142, p<0.001; advanced: n=6080, r=-0.136, p<0.001).

**Figure S4:**
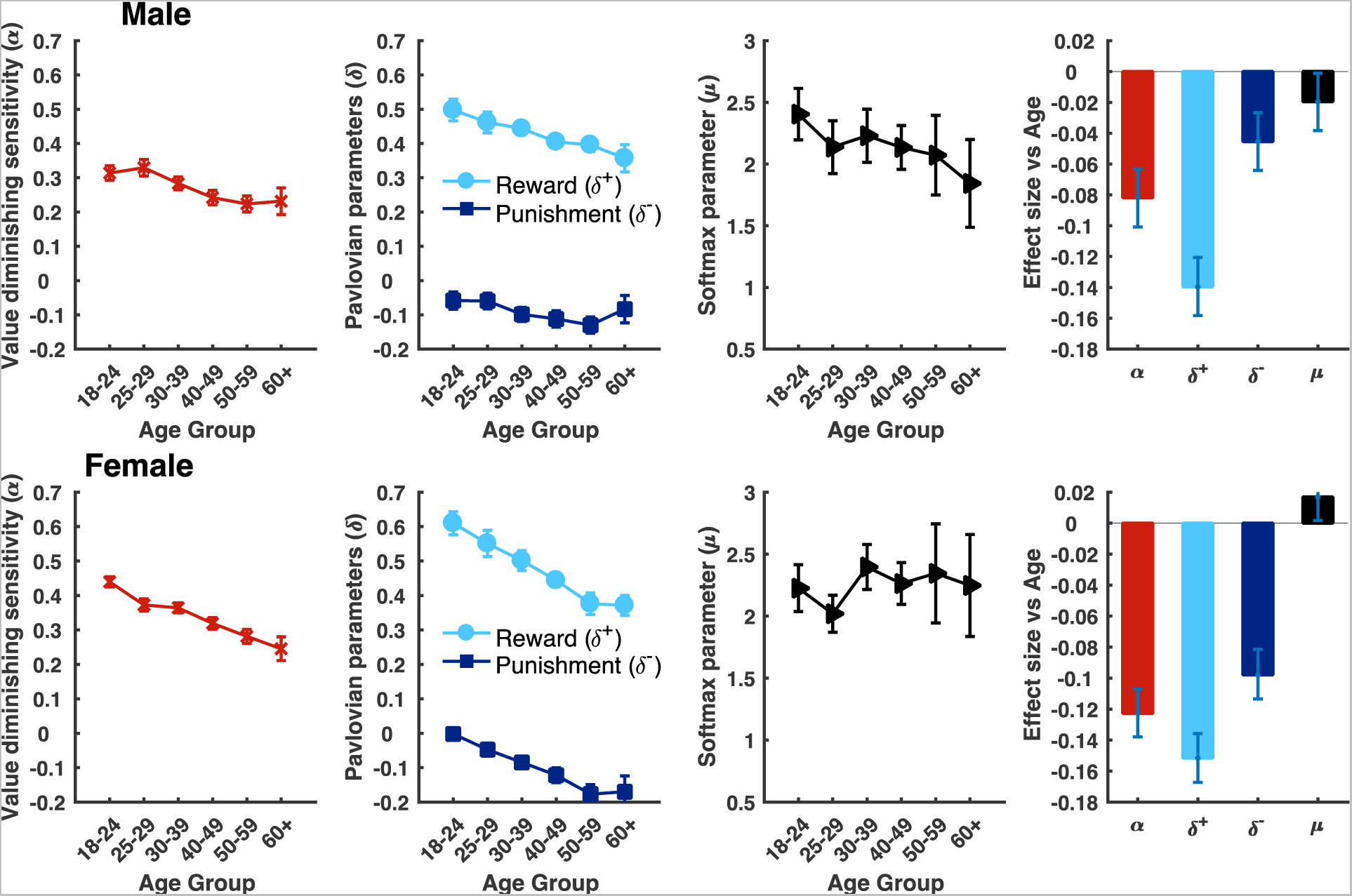
The change in approach-avoidance model parameters across the life span for each gender (top: male; bottom: female). Column 1 from left: α across age groups; Column 2: δ^-.^and δ^+.^across age groups; Column 3: μ across age groups; Column 4: age-related decline across the punishment and reward domain. The largest effect size was observed for the Pavlovian approach parameter (δ^+.^); Bars and error bars represent medians and bootstrapped 95%CIs.

**Figure S5:**
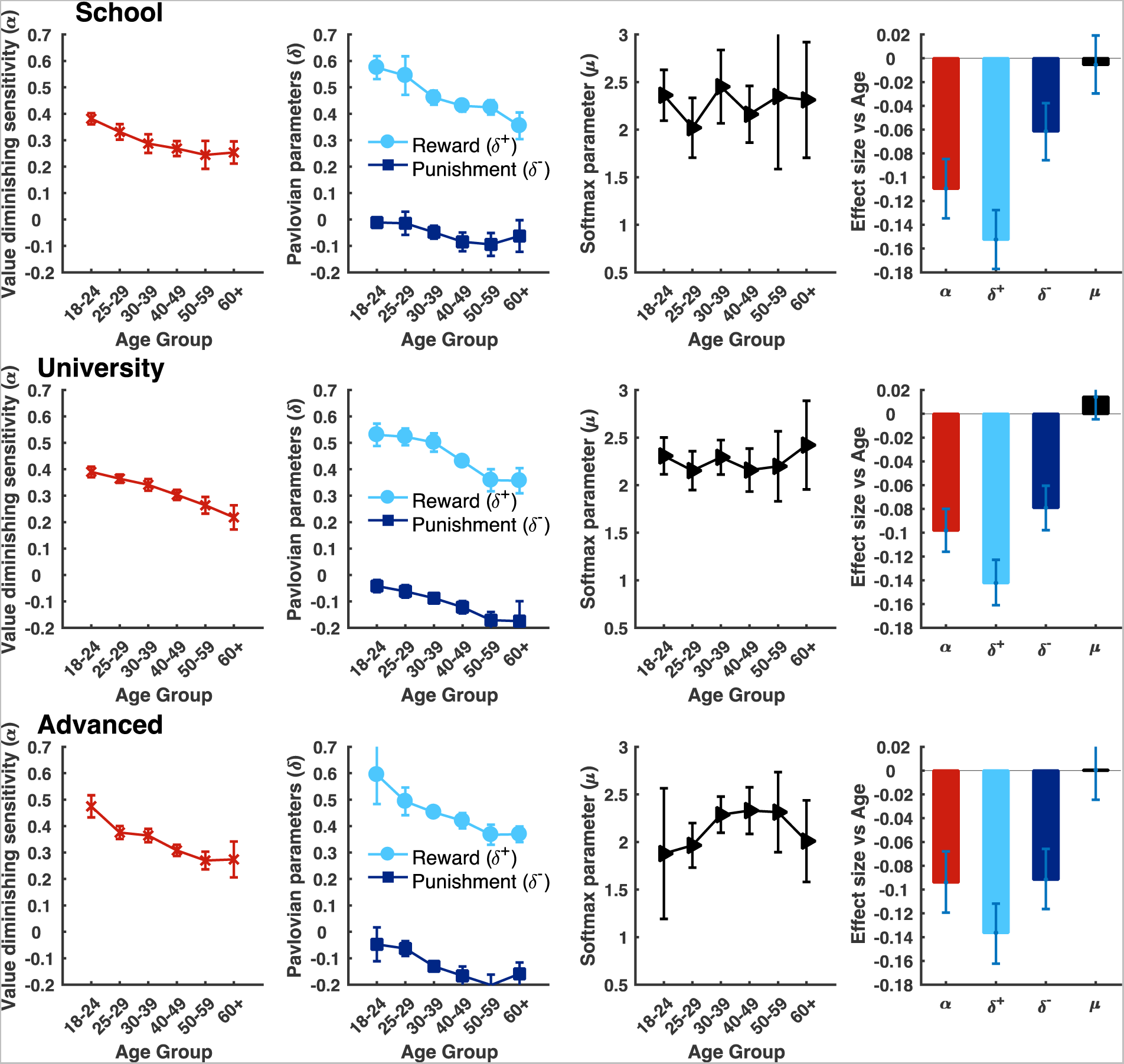
The change in approach-avoidance model parameters across the life span for each education level (top: school leavers; middle: university leavers; bottom: advanced (Masters, PhD). Column 1 from left: α across age groups; Column 2: δ^-^ and δ^+^ across age groups; Column 3: μ across age groups; Column 4: age-related decline across the punishment and reward domain. The largest effect size was observed for the Pavlovian approach parameter (δ^+^); Bars and error bars represent medians and bootstrapped 95%CIs.

### Correlation across the economic and motor decision-making tasks within each age group

Through the app-based platform a subset of participants (n=17,220) also performed an economic decision-making gambling task in which a similar approach-avoidance model was used to explain choice behaviour (see main text for relevant reference). Through correlation and median-split analysis we found a small but significant positive relationship for all main model parameters between the tasks. This relationship was relatively consistent across the lifespan whereby we found a positive correlation between these parameters within each age group (Figure S6 & S7). However, although the oldest age group (60+) showed a similar trend, we did not have enough power (participant numbers) to reliably detect effect sizes of 0.05 within this group. Specifically, whilst the 60+ age group (n=783) had 0.28 power to detect 0.05 effect sizes, the next oldest group (50-59, n=1541) had near double the amount of power of 0.5.

**Figure S6:**
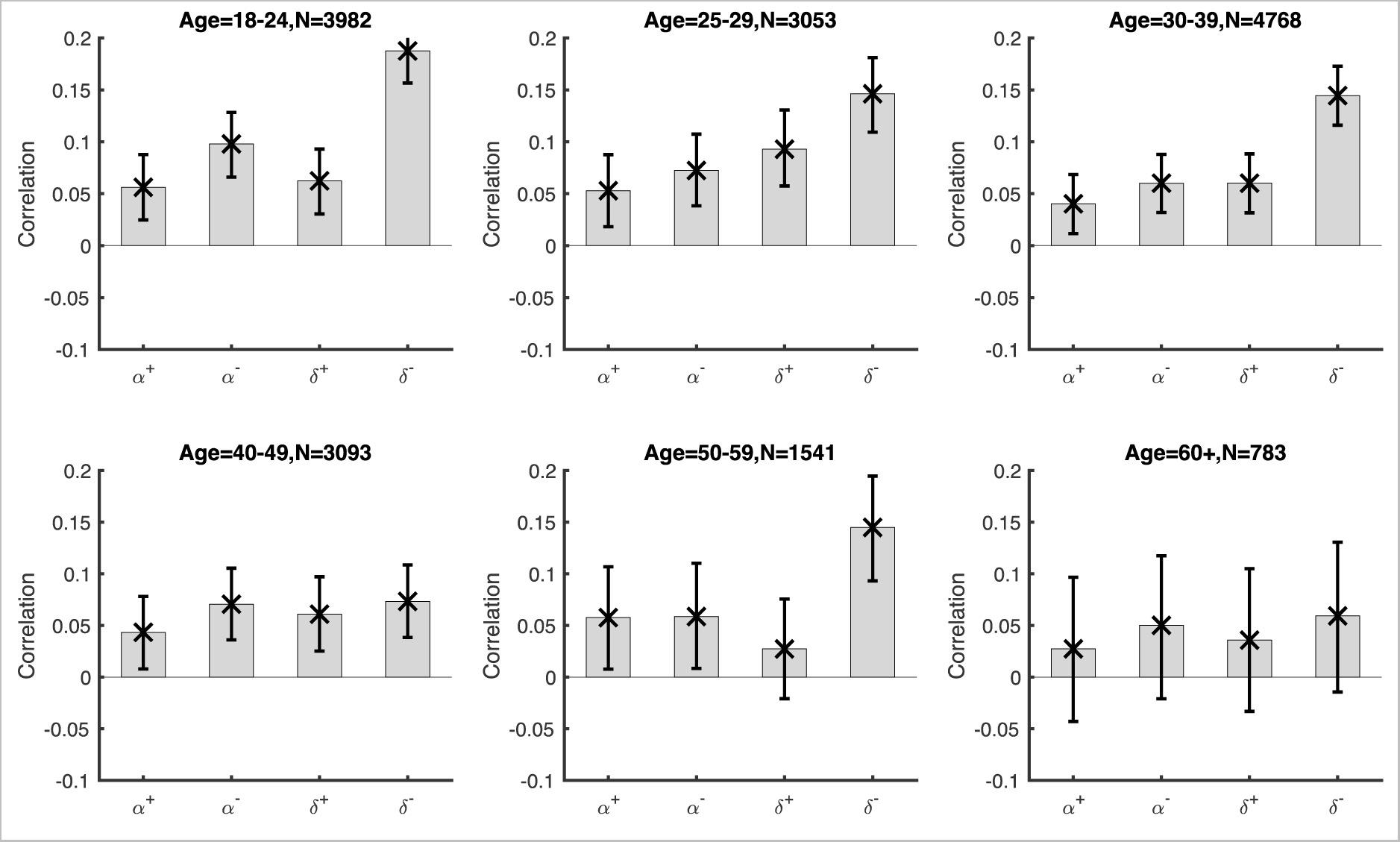
Correlation between motor and economic decision-making tasks for the main approach-avoidance model parameters within each age group. This relationship was relatively consistent across the lifespan whereby we found a positive correlation between these parameters within each age group. Note, the single α parameter of the motor decision-making model was correlated with both the α- and α+ parameters of the decision-making model. Error bars reflect bootstrapped 95% CIs.

**Figure S7:**
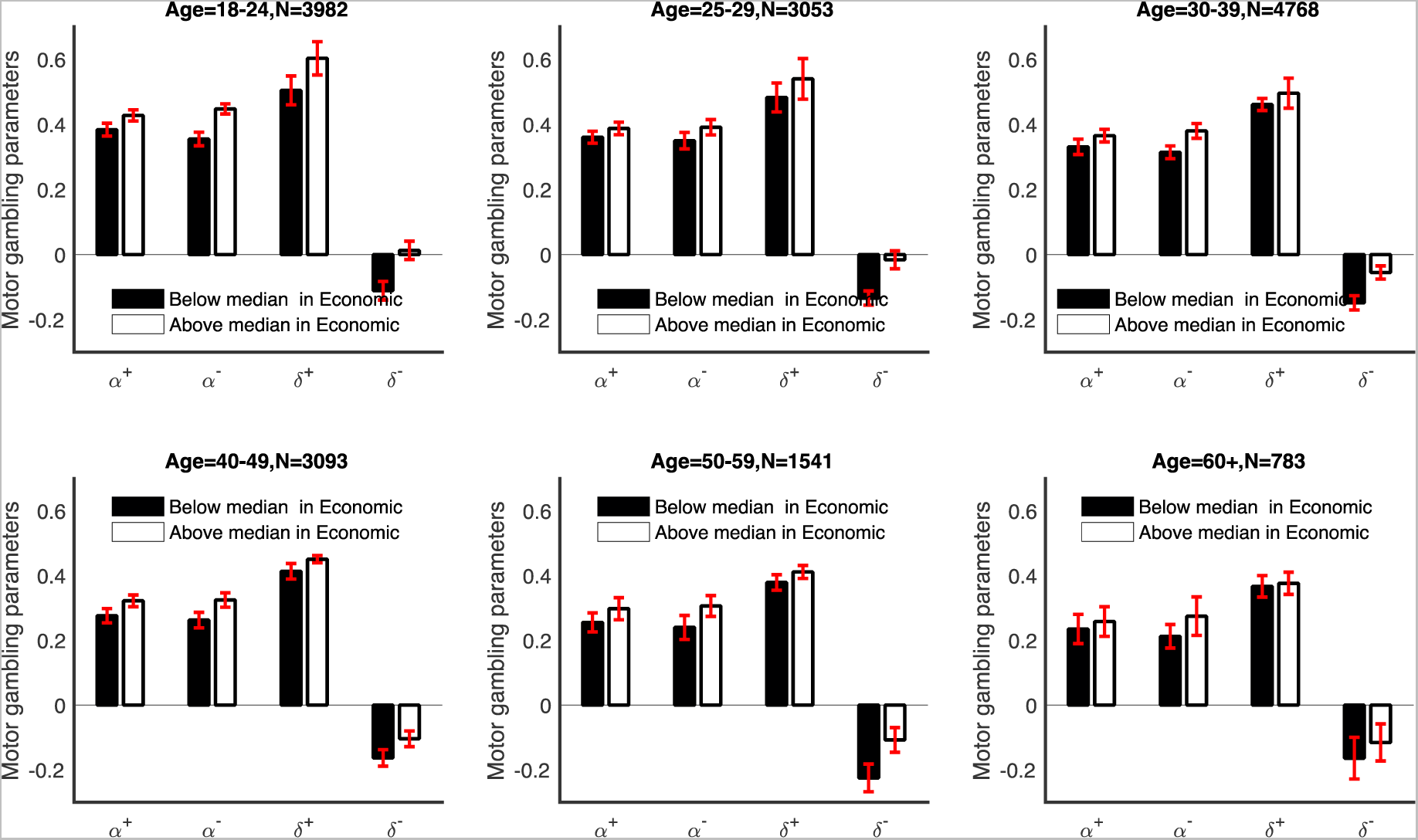
Motor decision-making approach-avoidance parameter values median split by economic parameter values within each age group. Filled bars denote participants with below-median values in the economic gambling task; Hollow bars for above-median. The participants with above-median risk parameters and Pavlovian parameters in the economic decision task had generally higher risk parameters and Pavlovian parameters in the motor gambling task. Bars/error bars reflect medians/bootstrapped 95%CIs.

### Model results when probability of success was based on each individual’s own performance

In the main results, the probability of success for a participant within a certain age group, using a certain screen size and facing a certain target size on each trial was estimated using the average success rate across all the participants with the same age, same screen size, and facing the same target size. Given the small amount of trials we had for each participant at each target size to estimate their probability of success, we believed this group average approach was the most valid estimate of success probability. However, Figure S8 shows that similar results are observed when probability of success is estimated based on each individual’s own data (i.e. the probability of success for a participant facing a certain target size was estimated using their own success rate over the same target size).

**Figure S8:**
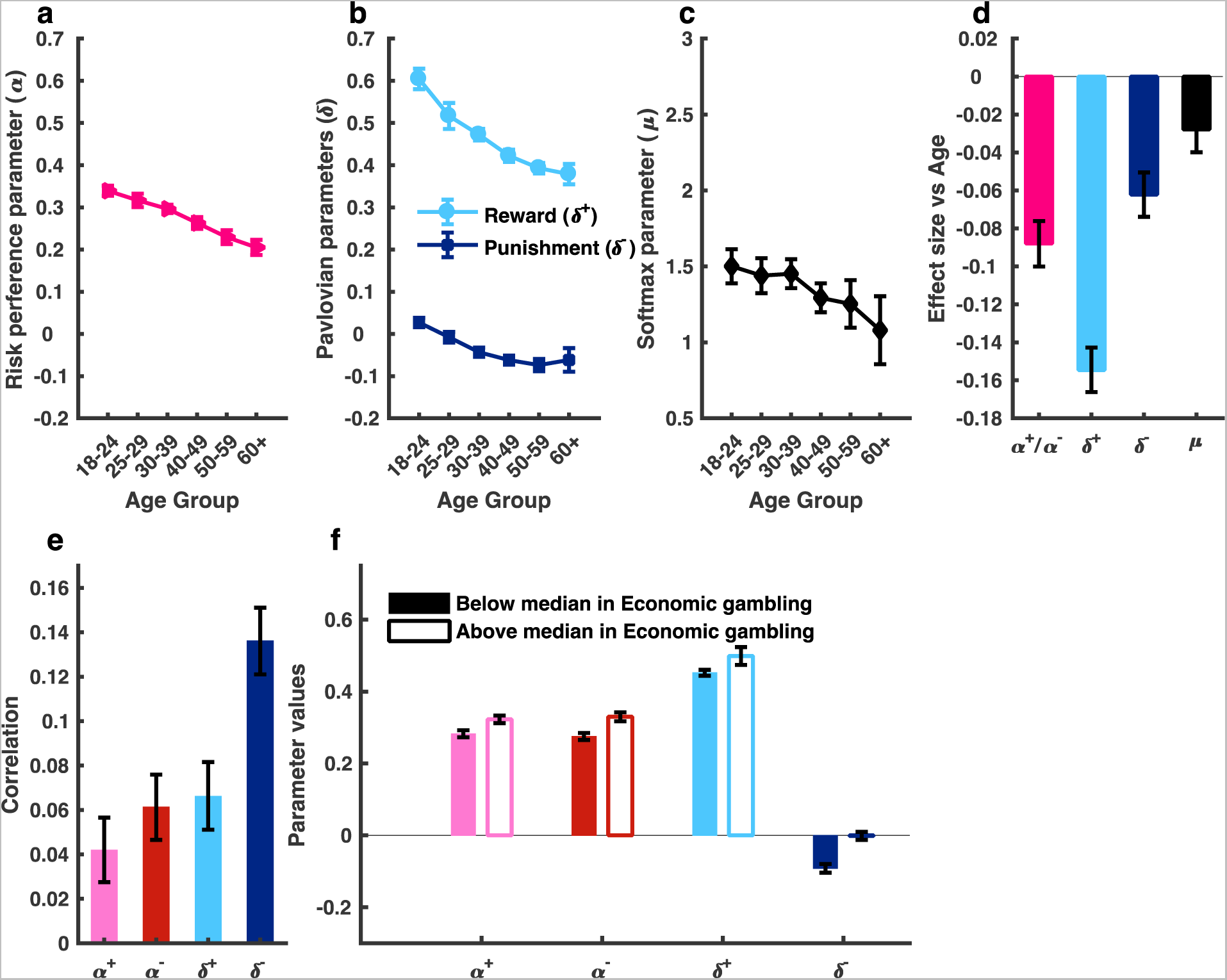
Model results when probability of success was based on each individual’s own performance (rather than group average). Similar results are observed when the probability of success was estimated based on each individual’s own data. (a) α across age groups; (b) δ^-^ and δ^+^ across age groups; (c) μ across age groups; (d) age-related decline across the loss and gain domain. The largest effect size was observed for the Pavlovian approach parameter (δ^+^); (e) positive correlation across motor and economic decision tasks for the main approach-avoidance model parameters; (f) median split. Filled bars denote participants with below-median values in the economic gambling task; Hollow bars for above-median. The participants with above-median risk parameters and Pavlovian parameters in the economic decision task had higher risk parameters and Pavlovian parameters in the motor gambling task. Bars/error bars reflect medians/bootstrapped 95%CIs.

## References

1. Valsecchi, M., Billino, J. & Gegenfurtner, K.R. Healthy Aging Is Associated With Decreased Risk-Taking in Motor Decision-Making. Journal of Experimental Psychology: Human Perception and Performance (2017).

2. Rutledge, R.B., et al. Risk Taking for Potential Reward Decreases across the Lifespan. Curr Biol 26, 1634–1639 (2016).

3. Tymula, A., Rosenberg Belmaker, L.A., Ruderman, L., Glimcher, P.W. & Levy, I. Like cognitive function, decision making across the life span shows profound age-related changes. Proc Natl Acad Sci U S A 110, 17143–17148 (2013).

4. Grubb, M.A., Tymula, A., Gilaie-Dotan, S., Glimcher, P.W. & Levy, I. Neuroanatomy accounts for age-related changes in risk preferences. Nat Commun 7, 13822 (2016).

5. Rutledge, R.B., Skandali, N., Dayan, P. & Dolan, R.J. Dopaminergic Modulation of Decision Making and Subjective Well-Being. J Neurosci 35, 9811–9822 (2015).

6. Rutledge, R.B., Skandali, N., Dayan, P. & Dolan, R.J. A computational and neural model of momentary subjective well-being. Proc Natl Acad Sci U S A 111, 12252–12257 (2014).

7. Wu, S.W., Delgado, M.R. & Maloney, L.T. Economic decision-making compared with an equivalent motor task. Proc Natl Acad Sci U S A 106, 6088–6093 (2009).

8. Trommershauser, J., Maloney, L.T. & Landy, M.S. Statistical decision theory and the selection of rapid, goal-directed movements. Journal of the Optical Society of America. A, Optics, image science, and vision 20, 1419–1433 (2003).

9. Trommershauser, J., Maloney, L.T. & Landy, M.S. Statistical decision theory and trade-offs in the control of motor response. Spatial vision 16, 255–275 (2003).

10. Jarvstad, A., Hahn, U., Rushton, S.K. & Warren, P.A. Perceptuo-motor, cognitive, and description-based decision-making seem equally good. Proc Natl Acad Sci U S A 110, 16271–16276 (2013).

11. Guitart-Masip, M., et al. Go and no-go learning in reward and punishment: interactions between affect and effect. Neuroimage 62, 154–166 (2012).

12. Guitart-Masip, M., Duzel, E., Dolan, R. & Dayan, P. Action versus valence in decision making. Trends Cogn Sci 18, 194–202 (2014).

13. Dayan, p., Niv, Y., Seymour, B. & Daw, N.D. The misbehavior of value and the discipline of the will. Neural Netw 19, 1153–1160 (2006).

14. Tversky, A. & Kahneman, D. Judgment under Uncertainty: Heuristics and Biases. Science 185, 1124–1131 (1974).

15. Kahneman, D. & Tversky, A. Prospect theory: an analysis of decision under risk. Econoemtrica 47, 263–291 (1979).

16. Sokol-Hessner, P., et al. Thinking like a trader selectively reduces individuals’ loss aversion. Proc Natl Acad Sci U S A 106, 5035–5040 (2009).

17. Frydman, C., Camerer, C., Bossaerts, P. & Rangel, A. MAOA-L carriers are better at making optimal financial decisions under risk. Proc Biol Sci 278, 2053–2059 (2011).

18. Hunt, L.T., Rutledge, R.B., Malalasekera, W.M., Kennerley, S.W. & Dolan, R.J. Approach-Induced Biases in Human Information Sampling. PLoS Biol 14, e2000638 (2016).

19. Trommershauser, J., Landy, M.S. & Maloney, L.T. Humans rapidly estimate expected gain in movement planning. Psychological science 17, 981–988 (2006).

20. Trommershauser, J., Gepshtein, S., Maloney, L.T., Landy, M.S. & Banks, M.S. Optimal compensation for changes in task-relevant movement variability. J Neurosci 25, 7169–7178 (2005).

21. Wu, S.W., Delgado, M.R. & Maloney, L.T. The neural correlates of subjective utility of monetary outcome and probability weight in economic and in motor decision under risk. J Neurosci 31, 8822–8831 (2011).

22. Jarvstad, A., Hahn, U., Warren, P.A. & Rushton, S.K. Are perceptuo-motor decisions really more optimal than cognitive decisions? Cognition 130, 397–416 (2014).

23. Freeman, S.M. & Aron, A.R. Withholding a Reward-driven Action: Studies of the Rise and Fall of Motor Activation and the Effect of Cognitive Depletion. J Cogn Neurosci 28, 237–251 (2016).

24. Freeman, S.M., Alvernaz, D., Tonnesen, A., Linderman, D. & Aron, A.R. Suppressing a motivationally-triggered action tendency engages a response control mechanism that prevents future provocation. Neuropsychologia 68, 218–231 (2015).

25. Freeman, S.M., Razhas, I. & Aron, A.R. Top-down response suppression mitigates action tendencies triggered by a motivating stimulus. Curr Biol 24, 212–216 (2014).

26. Takikawa, Y., Kawagoe, R., Itoh, H., Nakahara, H. & Hikosaka, O. Modulation of saccadic eye movements by predicted reward outcome. Exp Brain Res 142, 284–291 (2002).

27. Weiler, J.A., Bellebaum, C. & Daum, I. Aging affects acquisition and reversal of reward-based associative learning. Learning & memory 15, 190–197 (2008).

28. Mell, T., et al. Effect of aging on stimulus-reward association learning. Neuropsychologia 43, 554–563 (2005).

29. McNab, F. & Dolan, R.J. Dissociating distractor-filtering at encoding and during maintenance. J Exp Psychol Hum Percept Perform 40, 960–967 (2014).

30. Brown, H.R., et al. Crowdsourcing for cognitive science-the utility of smartphones. PLoS One 9, e100662 (2014).

31. von Neumann, J. & Morgenstem, O. Theory of Games and Economic Behavior (Princeton University Press, Princeton, 1944).

32. Mather, M., et al. Risk preferences and aging: the "certainty effect" in older adults' decision making. Psychology and aging 27, 801–816 (2012).

33. Boureau, Y.L. & Dayan, P. Opponency revisited: competition and cooperation between dopamine and serotonin. Neuropsychopharmacology 36, 74–97 (2011).

34. van Dyck, C.H., et al. Age-related decline in central serotonin transporter availability with [(123)I]beta-CIT SPECT. Neurobiology of aging 21, 497–501 (2000).

35. Backman, L., Nyberg, L., Lindenberger, U., Li, S.C. & Farde, L. The correlative triad among aging, dopamine, and cognition: current status and future prospects. Neurosci Biobehav Rev 30, 791–807 (2006).

36. Karrer, T.M., Josef, A.K., Mata, R., Morris, E.D. & Samanez-Larkin, G.R. Reduced dopamine receptors and transporters but not synthesis capacity in normal aging adults: a meta-analysis.

37. inger, B., Schuck, N.W., Nystrom, L.E. & Cohen, J.D. Reduced striatal responses to reward prediction errors in older compared with younger adults. J Neurosci 33, 9905–9912 (2013).

38. Swart, J.C., et al. Catecholaminergic challenge uncovers distinct Pavlovian and instrumental mechanisms of motivated (in)action. Elife 6(2017).

39. Chen, X., Mohr, K. & Galea, J.M. Predicting explorative motor learning using decision-making and motor noise. PLoS Comput Biol 13, e1005503 (2017).

40. >Gonzalez, R. & Wu, G. On the shape of the probability weighting function. Cogn Psychol 38, 129–166 (1999).

41. Akaike, H. A new look at the statistical model identification. IEEE transactions on automatic control 19, 716–723 (1974).

42. Schwarz, G. Estimating the dimension of a model. The annals of statistics 6, 461–464 (1978).

